# Exosome-mediated viral nucleic acid presentation in a crustacean reveals innate immunity evolution from a novel perspective

**DOI:** 10.1101/2022.11.08.515566

**Authors:** Yi Gong, Hang Hu, Xinshan Zhao, Weiqian Wei, Ming Zhang, Tran Ngoc Tuan, Hongyu Ma, Yueling Zhang, Kok-Gan Chan, Shengkang Li

## Abstract

As an enduring hot topic in the field of innate immunity, apoptosis is widely considered as an effective approach to eliminate pathogenic microbes, and possesses crucial role during host-pathogen interactions. Recently, researchers have found that virus-containing host cells could transmit apoptotic signals to surrounding uninfected cells during infection, but the mechanism remains unclear. Here, we found that exosomes secreted by WSSV-infected mud crab hemocytes contain viral nucleic acid wsv277, which could be transported to the recipient cells and further expressed viral protein with phosphokinase activity. Besides, by using transcriptome, proteome, ChIP-seq and coIP techniques, the results revealed that wsv277 could activate the transcription and translation of apoptotic genes via interacting with CBF and EF-1α, so as to suppress the spread of virus infection by inducing apoptosis of the surrounding cells. Therefore, for the first time, our study proved that the components of DNA virus could be encapsulated into exosomes, and elucidated the mechanism of apoptotic signal transduction between cells from the perspective of exosome. Moreover, comparing those of invertebrates, we hypothesized that the innate immune system of vertebrate was weakened or degenerated, allowing exosomes to be employed by virus as the vehicle for immune escape.

**Significance statement:** Our study revealed that the components of DNA virus could be packaged and transmitted through the exosomes of lower invertebrates, which strongly demonstrated the diversity of exosome-mediated viral immunity and its universality in animals. Furthermore, we elucidated the mechanism of apoptotic signal transduction between cells from the perspective of exosome, and revealed a novel strategy for the host to cope with viral infection. Since exosome is an important innate immune regulation approach, we hypothesize that the innate immune system of vertebrates is weakened or degenerated, allowing exosomes to be employed by virus as the vehicle for immune escape.

## Introduction

Generally, the immune systems in animals are often classified into highly specific antibody-based adaptive immune response and nonspecific innate immune response^1^. The innate immune system, which uses evolutionarily conserved pattern recognition receptor systems^2^, including TLRs (Toll-like receptors), CLRs (C-type Lectin receptors), NLRs (Nod-like receptors) and RLRs (RIG-1 like receptors) to activate downstream cellular and humoral immune effectors^3^, is the host’ s first line of defense against pathogens and the main approach for invertebrates to resist harmful microbes^4^ Due to the lack of immunoglobulin, the humoral immunity in invertebrates relies only on a series of enzymes and molecules in the hemolymph^5^, such as phenoloxidase, lectin, hemocyanin and antimicrobial peptides^6^. While the cellular immunity in invertebrates is mainly accomplished by hemolymph cells through phagocytosis, encapsulation, aggregation and nodule formation^7^ In recent years, with the deepening of the research on innate immune regulation, many novel regulatory approaches have been found continuously^8,9^ Among them, the immunological role of exosomes during pathogen-host interactions is becoming the research hotspot of innate immunity^10^.

Exosomes are nanosized extracellular vesicles measuring 30-120 nm in diameter, which can be produced by almost all types of cells under physiological and pathological conditions^11^. As a form of intercellular vesicular transport, exosomes can be secreted by donor cells and transferred to target cells through cytomembrane fusion^12,13^, and serve as intercellular communication mediators by packaging bio-cargoes, such as lipids, proteins and nucleic acids^14,15^. Specific membrane proteins that highly enriched in exosomes are often used as indicators for the identification of exosomes, such as TSG101, CD9, CD63 and CD81^16^. Recently, the immunological role of extracellular vesicles in pathogen-host interactions have been extensively investigated^17^, numerous studies have pointed out the importance of exosomes during viral pathogenesis and host immune responses^10^. During virus infection, it is believed that the infected host cells are capable of excreting exosomes containing viral or host component to nearby cells, which further help to modulate host innate immune response and possess impact on the fate of the infection process^18,19^ Therefore, in-depth exploration of the biological function of exosomes is of great significance to enrich the regulatory mechanism of innate immunity.

As ideal creatures for exosome to exert biological functions, crustaceans possess an open circulatory system, where hemocytes, nutrients, hormones and oxygen circulate together in the hemolymph^20^, this type of circulatory system enables exosomes to be transported to almost all tissues and organs of the body through hemolymph flow^21^. More importantly, unlike vertebrates, crustaceans belong to invertebrates and mainly rely on the innate immune system to resist pathogenic microorganism infection^22^, which can effectively eliminate the impact of acquired immunity on the infection process^23^. Therefore, based on the above considerations, crustaceans are more suitable biosystem for exosomes to perform innate immune-related functions compared with vertebrates. However, as a topical research area, exosome-associated studies are largely focused on higher organisms^24^ In invertebrates, the immunological functions of exosomes have been reported only in *Drosophila* and mud crab, and were found to be involved in the regulation of viral and bacterial infection by mediating hemolymph microbiota homeostasis and apoptosis^19,25,26^. But overall, research about how exosomes participate in the antiviral innate immune response of invertebrates is still in its infancy stage, relevant theoretical and experimental support is urgently needed.

In an attempt to comprehensively reveal the mechanism of exosomes in regulating host immune response and the impact on the fate of viral infection process, WSSV (White spot syndrome virus)-infected mud crab *Scylla paramamosain* was used as the research model in this study. Herein, we found that five viral mRNAs were specifically packaged by exosomes during the infection process. Among them, wsv277 could be delivered to surrounding exosome-receiving cells and translated into viral proteins, and further mediated the transcription and translation of apoptotic genes through interacting with CBF (CCAAT-binding factor) and EF-1α (Elongation factor 1 alpha), which eventually suppressed virus invasion by inducing apoptosis of neighboring cells.

## Results

### Screening of viral nucleic acids in the exosomes of WSSV-infected mud crab

In order to determine whether exosomes secreted by invertebrate are capable of transporting viral genomic nucleic acids like mammals, we tried to analyze the nucleic acid composition in the exosomes of mud crabs before/ after WSSV infection. The typical cup-shaped structure of exosomes collected from the hemolymph of WSSV-injected (exosome-WSSV) and PBS-injected (exosome-PBS) mud crabs were observed under TEM (Transmission electron microscopy) (Fig. 1A). Besides, the quantities and sizes of the isolated exosomes were measured by nanoparticle tracking analysis (NTA) (Fig. 1B), and the purity were further ascertained via detecting the exosomal markers CD63, CD9, TSG101 and the cytoplasmic marker Calnexin (Fig. 1C). The data indicate successful isolation of exosomes from mud crab challenged with WSSV or PBS.

**Fig 1.**
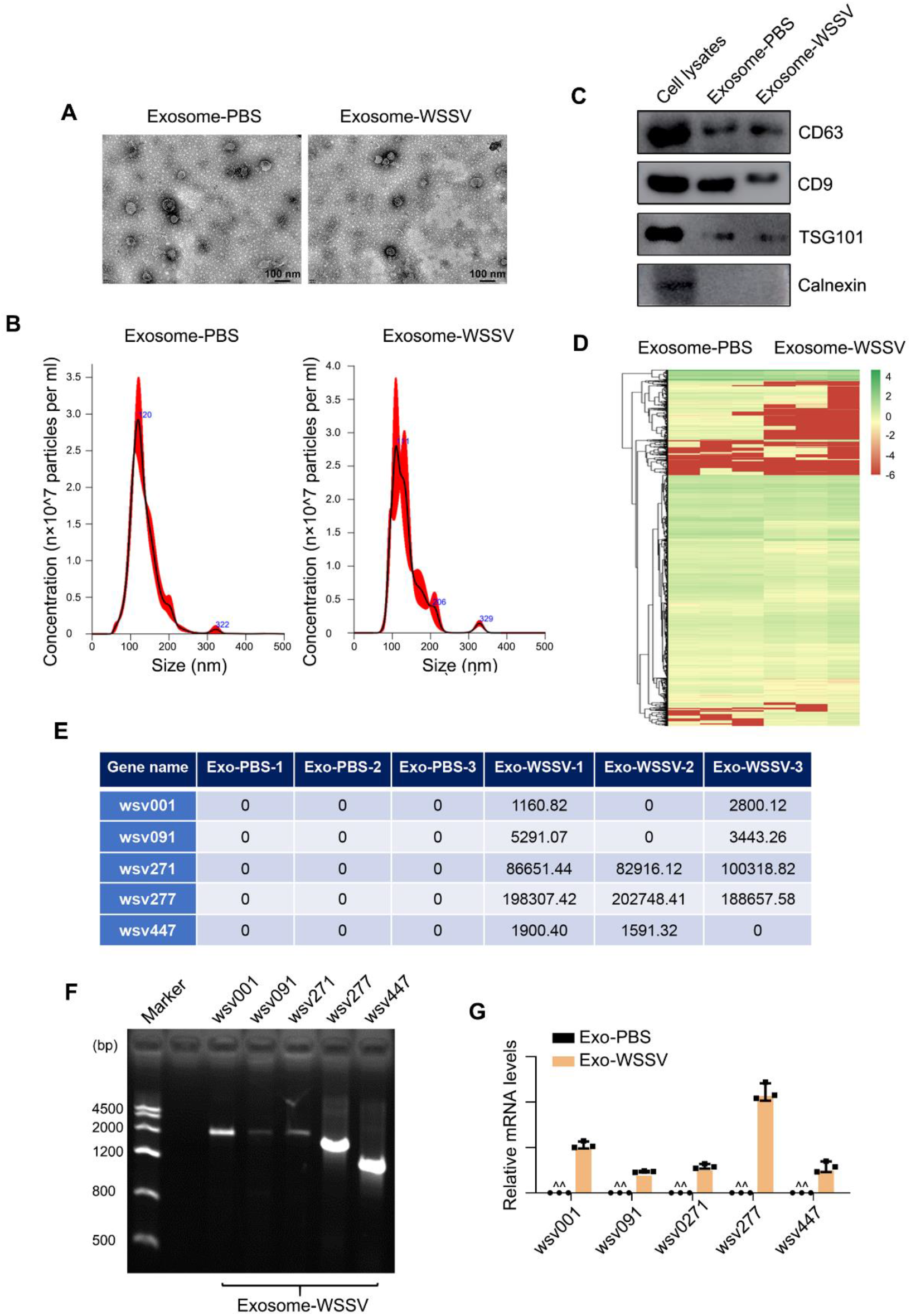
Screening of viral nucleic acids in exosomes. **(A-B)** Exosomes isolated from mud crabs with specified treatments (PBS and WSSV injection for 48 h) were detected by TEM **(A)** and NTA **(B)**. Scale bar, 100 nm. **(C)** Western blot detection of exosomal marker proteins (CD9, CD63 and TSG101) and cytoplasmic marker protein Calnexin in cell lysates and exosomes. (**D-E**) Transcriptome sequencing of the above isolated exosomes. Exosomal mRNAs were presented in a heatmap **(D)**, five WSSV nucleic acids were identified from the sequencing data through blasting with WSSV genome **(E)**. **(F-G)** Detection of viral nucleic acids in the exosomes. Exosomes were isolated from mud crabs challenged with WSSV for 48 h, followed by PCR **(F)** and qPCR **(G)** assay, exosome-PBS group was served as negative control, the brightness of nucleic acid band in agarose gel did not represent the expression levels. All data represented were the mean ± s.d. of at least three independent experiments (**, *P*<0.01).

Next, transcriptome sequencing was performed on the above isolated exosomes (Fig. 1D), and 5 WSSV-specific nucleic acids (wsv001, wsv091, wsv271, wsv277 and wsv447) were identified in exosome-WSSV group via blasting with WSSV genome (Fig. 1E). Then, PCR and qPCR were both conducted to confirm the existence of the viral nucleic acids (Fig. 1F and 1G). Taken together, the above findings revealed that 5 viral mRNAs were specific encapsulated in the exosomes of mud crab during WSSV infection, and the level of wsv277 was the highest among them.

### wsv277-containing exosomes suppress WSSV replication via apoptosis

Our previous study showed that exosome secreted from WSSV-challenged mud crab could inhibit virus infection by inducing apoptosis^19^, based on this consideration, we tried to explore whether this process was mediated by the viral mRNAs packaged in exosomes. Through co-injection of exosomes and specific viral siRNAs (Fig. S1), the WSSV copy number in WSSV+Exo-WSSV+wsv277-siRNA group was significantly increased compared with control (Fig. 2A), indicating the essential role of wsv277 in exosome-mediated virus suppression. To further confirm the results, we detected the involvement of wsv277 in exosome-mediated apoptosis induction. The data of annexin V, caspase 3 activity and mitochondrial membrane potential analysis showed that wsv277 was required for exosome-mediated apoptosis (Fig. 2B–2D). Besides, the mRNA and protein levels of apoptotic gene BOK and anti-apoptotic gene ICAD also showed the same trends (Fig. 2E–2F). Moreover, to reveal whether wsv277-mediated virus suppression was relevant to apoptosis, apoptosis inducer MG-132 was co-injected with Exo-WSSV and wsv277-siRNA, the results showed that MG-132 significantly inhibited the increase of viral load induced by wsv-277 interference (Fig. 2G). Taken together, these data indicated that exosome-mediated apoptosis and virus suppression in mud crab were regulated by viral mRNA wsv277 packaged by exosomes.

**Fig 2.**
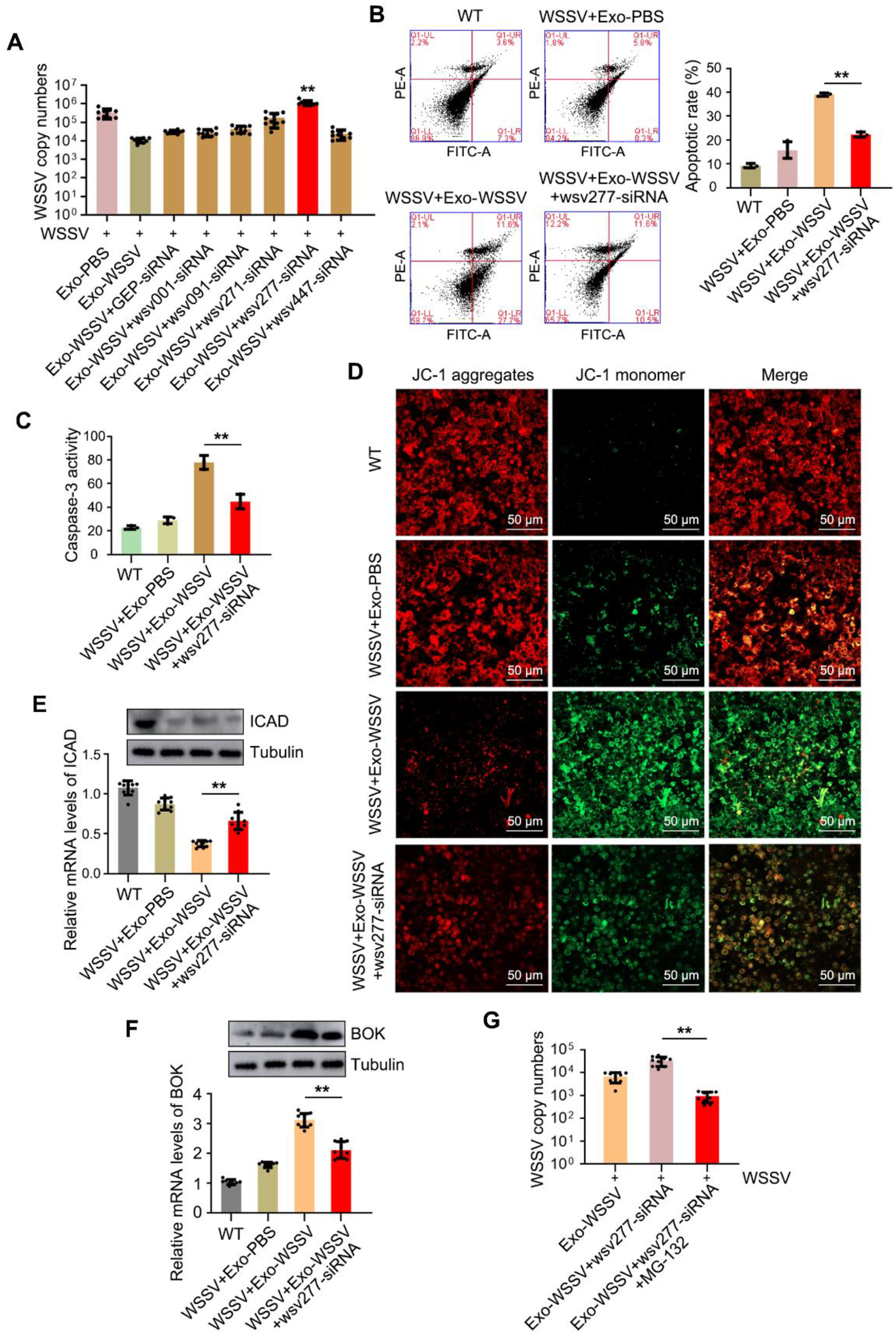
wsv277 contained in exosomes could inhibit virus infection by inducing apoptosis. **(A)** The effects of exosome-containing viral nucleic acids on exosome-mediated virus suppression. siRNAs targeting wsv001, wsv091, wsv271, wsv277 and wsv447 were co-injected with exosomes into WSSV-challenged mud crabs, followed by viral copy numbers detection via qPCR. **(B-F)** The influence of wsv277 on exosome-mediated apoptosis induction. The isolated exosomes were co-injected with WSSV into mud crabs for 48 h, followed by wsv277-siRNA treatment, then the apoptotic levels of the hemocytes were examined through annexin V analysis **(B)**, caspase 3 activity assay **(C)**, mitochondrial membrane potential measurement **(D)**, anti-apoptotic (ICAD) **(E)** and apoptotic (BOK) **(F)** genes’ detection, respectively. Untreated mud crabs were used as negative control. **(G)** The involvement of apoptosis induction during exosomal wsv277-mediated virus suppression. Mud crabs were co-injected with exosomes, WSSV, wsv277-siRNA and apoptosis inducer MG-132, then virus copy numbers were detected. All the data were the average from at least three independent experiments, mean ± s.d. (**, *P*<0.01).

### wsv277 mRNA is translated into a phosphokinase in exosome-recipient cells

To explore the regulatory mechanisms of wsv277 in exosome-mediated antiviral immune response, the key question is to determine whether wsv277 can be transmitted into crab cells along with exosomes. Thus, we injected exosomes into mud crabs and then detected the existence of wsv277 in hemocytes. The results of both agarose gel electrophoresis and fluorescence *in situ* hybridization (FISH) showed that wsv277 mRNA could be transported into cells and localized in the cytoplasm (Fig. 3A and 3B). Furthermore, through western blot and immunofluorescence analysis, it was found that wsv277 mRNA packaged by exosomes could be translated into viral protein in hemocytes (Fig. 3C and 3D).

**Fig 3.**
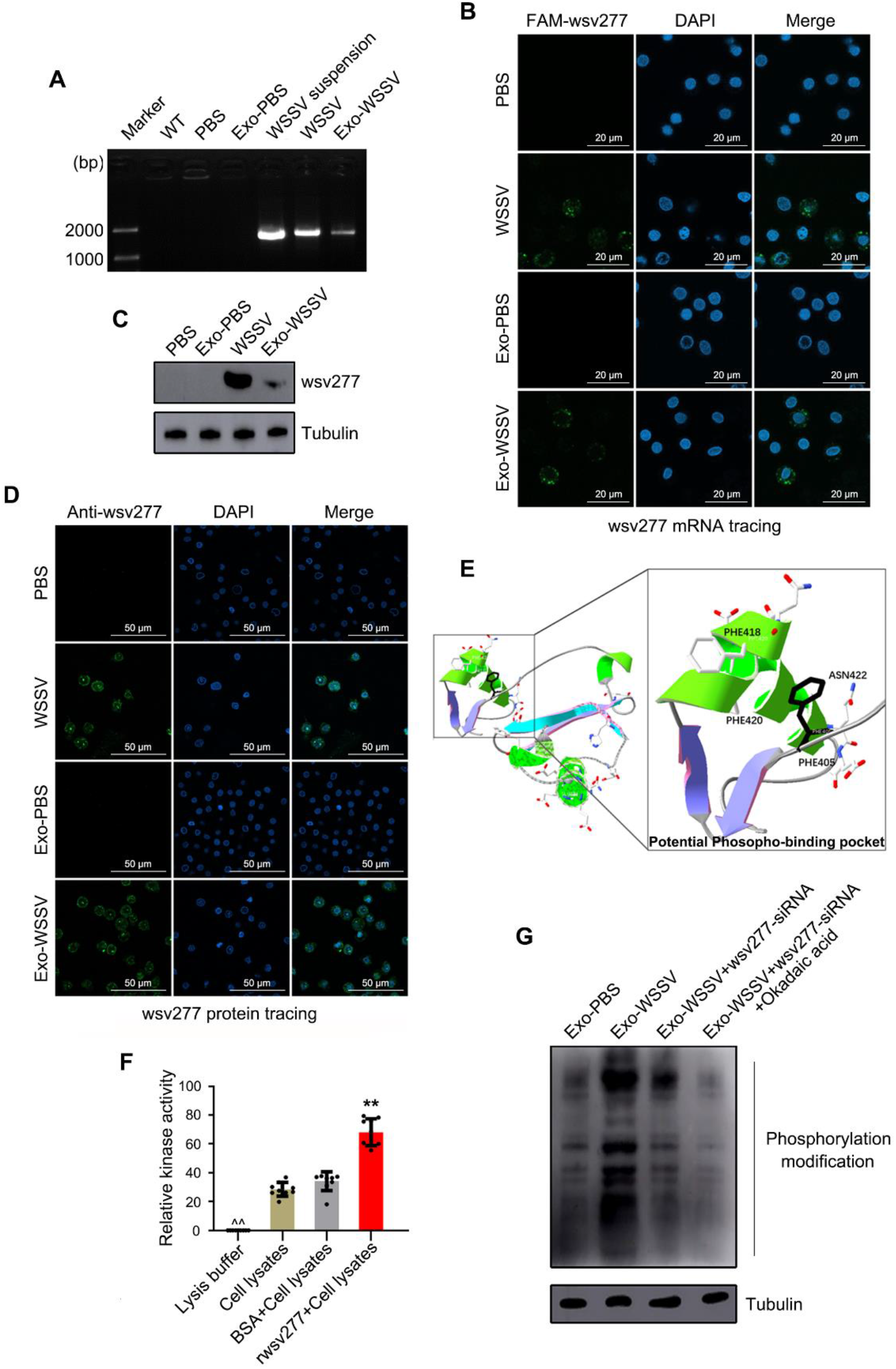
wsv277 mRNA is transmitted to exosome-recipient cells and translated into a phosphokinase. **(A-B)** wsv277 mRNA is transmitted to exosome-recipient cells. Exo-PBS, Exo-WSSV and WSSV was injected into mud crabs, respectively, then the ORF region of wsv277 was amplified by PCR, followed by agarose gel electrophoresis, WSSV suspension was used as positive control **(A)**; The localization of wsv277 mRNA in mud crab hemocytes was detected with FAM-labeled wsv277 mRNA probe (green), the nucleus was stained with DAPI (blue). Scale bar, 20 μm **(B)**. **(C-D)** wsv277 mRNA packaged in exosomes was translated into a viral protein in mud crab hemocytes. Exo-PBS, Exo-WSSV and WSSV was injected into mud crabs, respectively, then wsv277 was detected by western blot assay, tubulin was used as internal reference **(C)**; The localization of wsv277 protein was detected by immunofluorescence using wsv277-specific antibody. Scale bar, 50 μm **(D)**. **(E)** Tertiary structure prediction of wsv277 protein. Amino acids composed of PHE405, PHE418, PHE420 and ASN422 possess potential phosphokinase activity. **(F)** Kinase activity detection of recombinant wsv277 protein *in vitro*, BSA protein was used as negative control. **(G)** The effects of exosomes and wsv277 on protein phosphorylation in mud crab. Okadaic acid was used as a kind of phosphatase inhibitor. Experiments were performed at least in triplicate and the data represented were the mean ± s.d. (**, *P*<0.01).

Then, to reveal the potential functions of the translated wsv277 protein, the amino acid sequence of wsv277 was subjected to domain prediction by bioinformatics tools, and phosphorus residues were found on PHE405, PHE418, PHE420 and ASN422 in the tertiary structure (Fig. 3E), indicating that wsv277 protein may possess phosphokinase activity. To confirm this conjecture, we detected the phosphorylation ability of *r*wsv277 (recombinant wsv277 protein) to hemocyte lysates, and found that *r*wsv277 was capable to phosphorylate hemocyte lysates *in vitro* (Fig. 3E). In addition, we co-injected exosomes and wsv277-siRNAs into mud crabs and then measured the phosphorylation levels of hemocyte proteins, the results showed that Exo-WSSV could significantly increase the phosphorylation levels of hemocyte proteins, whereas this process was suppressed when wsv277 was silenced (Fig. 3F). Taken together, these findings suggested that wsv277 mRNA encapsulated in exosomes could be transmitted and translated into viral protein with phosphokinase activity in exosome-recipient cells.

### Screening and validation of wsv277-interacting proteins

Kinases are generally considered to be tightly relevant to the regulation of gene expression^27^, for this reason, we performed transcriptome analysis on the hemocytes of mud crab challenged with exosomes and wsv277-siRNA (Fig. 4A). Among the differentially expressed genes, 428 genes were found to be altered in Exo-WSSV group compared with Exo-PBS, and with no significant difference when wsv277 was silenced (Fig. S2A), indicating that wsv277 was required for the transcription of these genes. Besides, KEGG and GO analysis showed that many of wsv277-regulated genes were involved in apoptosis process (Fig. 4B and S2B), which was consistent with the above data. Next, combined with GST pull-down and mass spectrometry, we obtained a series of potential wsv277-interacting proteins in mud crab (Fig. 4C and 4D). Among them, transcription factor CBF and translation elongation factor EF-1α were selected since they possess gene expression regulatory function. Then, immunoprecipitation analysis performed with hemocytes lysates revealed that wsv277 could form complex with CBF and EF-1 α, respectively, while no direct interaction was existed between CBF and EF-1α (Fig. 4E). Besides, the above results were further confirmed by coIP experiment conducted in *Drosophila* S2 cells (Fig. 4F–4K). Taken together, these data demonstrated that wsv277 could form complex with CBF and EF-1 α, respectively, which possess the ability to modulate gene expressions.

**Fig 4.**
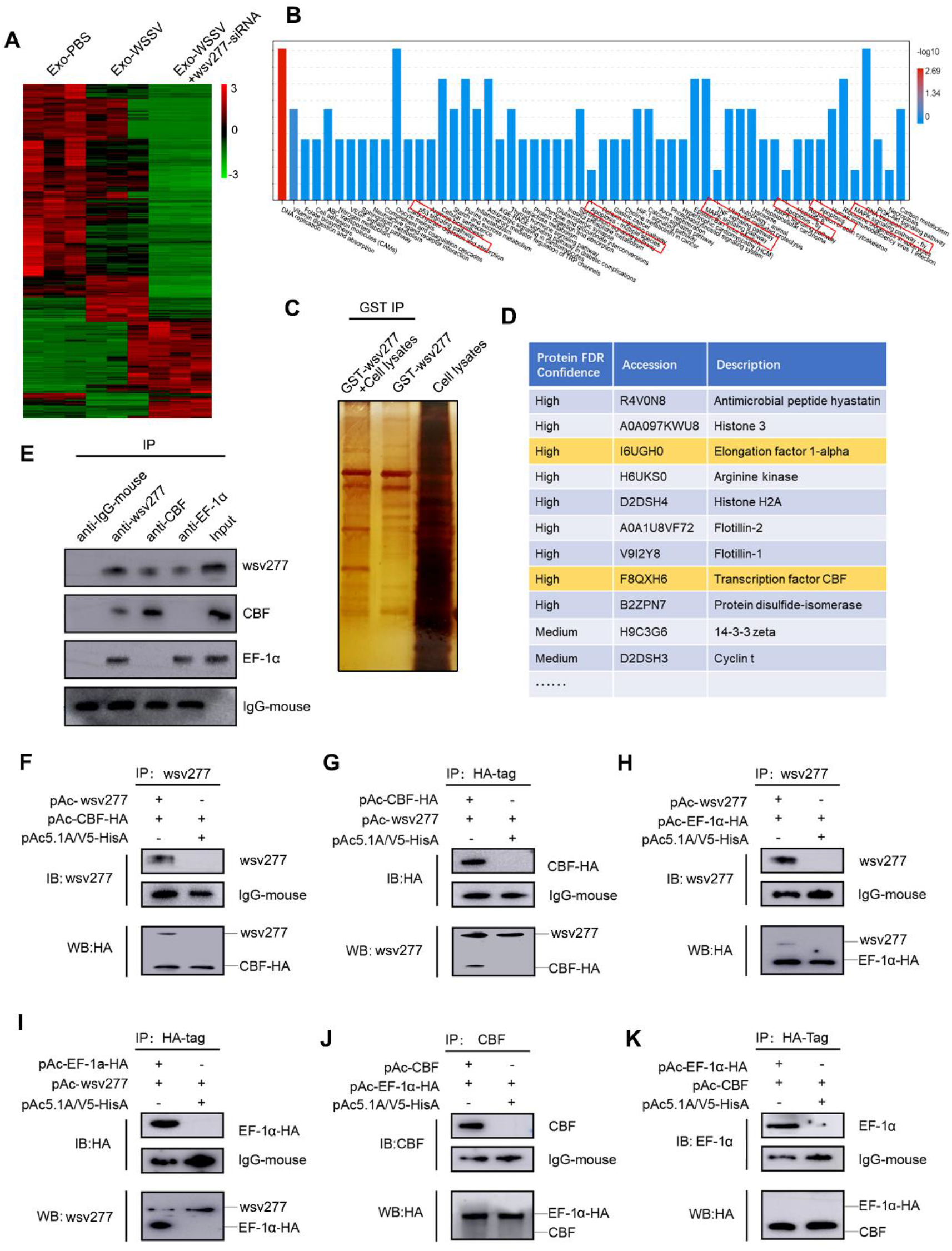
Screening and validation of wsv277-interacting proteins. **(A)** Transcriptome sequencing of mud crab hemocytes after treated with exosomes and wsv277-siRNA. 92366 differential expressed genes were presented in heatmap. **(B)** KEGG annotation of 428 differential expressed genes shown in Fig. S2A. The apoptosis-related pathways in the GO histogram were marked by red boxes. **(C-D)** Identification of proteins bound to wsv277. The cell lysates of hemocytes were incubated with recombinant GST-labeled wsv277 protein, followed by separated with SDS-PAGE and silver staining **(C)**; and identified by mass spectrometry **(D)**. **(E)** The interactions between wsv277, CBF and EF-1α in mud crab. Cell lysates were subjected to immunoprecipitation analysis with anti-wsv277 IgG, anti-CBF IgG and anti-EF-1α IgG, followed by western blot analysis using the indicated antibodies. **(F-G)** coIP analysis between wsv277 and CBF in S2 cells. wsv277 and/or HA-tagged CBF were co-transfected into S2 cells and then subjected to anti-wsv277 or anti-HA immunoprecipitation. **(H-I)** coIP analysis between wsv277 and EF-1α in S2 cells. **(F-G)** coIP analysis between CBF and EF-1α in S2 cells.

### CBF and EF-1α are required for exosome-mediated anti-viral immune response

To clarify whether CBF and EF-1 α were involved in exosome-mediated apoptosis and WSSV suppression, we knocked down the expression of CBF and EF-1α and then detected their influence on apoptosis (Fig. S3A–S3D). The results of annexin V, caspase 3 activity and mitochondrial membrane potential analysis showed that both CBF and EF-1α possess positive influence on apoptosis (Fig. 5A–5C). Besides, the expression of CBF and EF-1α were significantly increased during WSSV infection (Fig. 5D), while silencing of CBF and EF-1α promoted virus replication (Fig. 5E), indicating that both CBF and EF-1α could suppress WSSV infection. Next, to reveal the roles of CBF and EF-1 α during exosome-mediated anti-viral immune response, exosomes and siRNAs for CBF and EF-1α were co-injected into WSSV-challenged mud crabs, and the data of annexin V, caspase 3 activity and mitochondrial membrane potential analysis showed that silencing of CBF and EF-1α remarkably decreased apoptosis levels compared with control group (Fig. 5F–5H). In addition, through detecting the copy number of virus particles, we found that CBF and EF-1α were required for exosome-mediated WSSV suppression (Fig. 5I). Taken together, these finding suggested that CBF and EF-1α were the downstream targets for exosome-induced apoptosis and subsequent suppression of viral replication.

**Fig 5.**
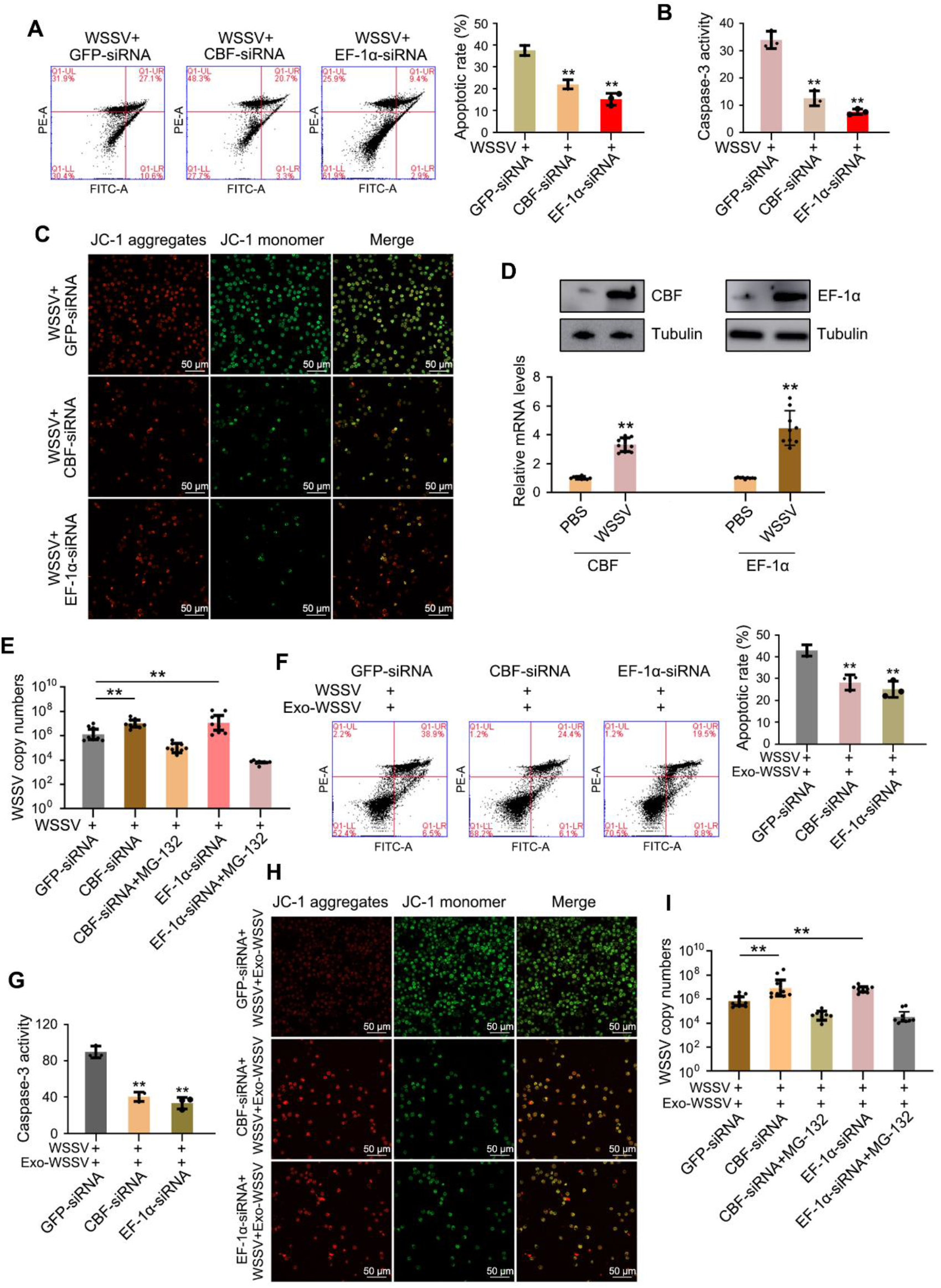
CBF and EF-1α contribute to exosome-mediated apoptosis and virus suppression. (**A-C**) The effects CBF and EF-1α on apoptosis during WSSV infection. Mud crabs were treated with WSSV and siRNAs targeting CBF or EF-1α for 48 h, then the apoptotic levels of the hemocytes were examined through annexin V analysis **(A)**, caspase 3 activity detection **(B)** and mitochondrial membrane potential measurement **(C)**. **(D)** The expression levels of CBF and EF-1α during WSSV infection in mud crab. Mud crabs were challenged to WSSV infection for 48 h, then the mRNA and protein levels of CBF and EF-1α were detected through qPCR and western blot, respectively. **(E)** The influence of CBF and EF-1α on WSSV infection. Mud crabs were cotreated with WSSV and siRNAs targeting CBF or EF-1α, followed by viral copy numbers detection via qPCR. **(F-H)** The influence of CBF and EF-1α on exosome-mediated apoptosis induction. The isolated exosomes were co-injected with WSSV into mud crabs for 48 h, followed by CBF/ EF-1α-siRNA treatment, then the apoptotic levels of the hemocytes were examined through annexin V analysis **(F)**, caspase 3 activity assay **(G)**, mitochondrial membrane potential measurement **(H). (I)** The involvement of CBF and EF-1α during exosome-mediated virus suppression. Mud crabs were co-injected with exosomes, WSSV and CBF/ EF-1α-siRNA, then virus copy number was detected. The results are based on three parallel experiments and shown as mean values ± s.d. (*, *P*<0.05, **, *P*<0.01).

### Underlying mechanisms of wsv277-CBF complex-mediated apoptosis

To reveal the mechanism and role of CBF in exosome-mediated apoptosis, we tried to analyze the phosphorylation levels of CBF after exosomes treatment, since the above findings indicated that wsv277 protein possess phosphokinase activity. The results showed that Exo-WSSV could phosphorylate CBF at tyrosine sites by packaging wsv277 (Fig. 6A). Then, the influence of the phosphorylation modification on the cellular localization of CBF was detected, the results of both western blot analysis and immune-fluorescence detection revealed that CBF would be translocated to the cell nucleus after phosphorylation by wsv277 (Fig. 6B and S4).

**Fig 6.**
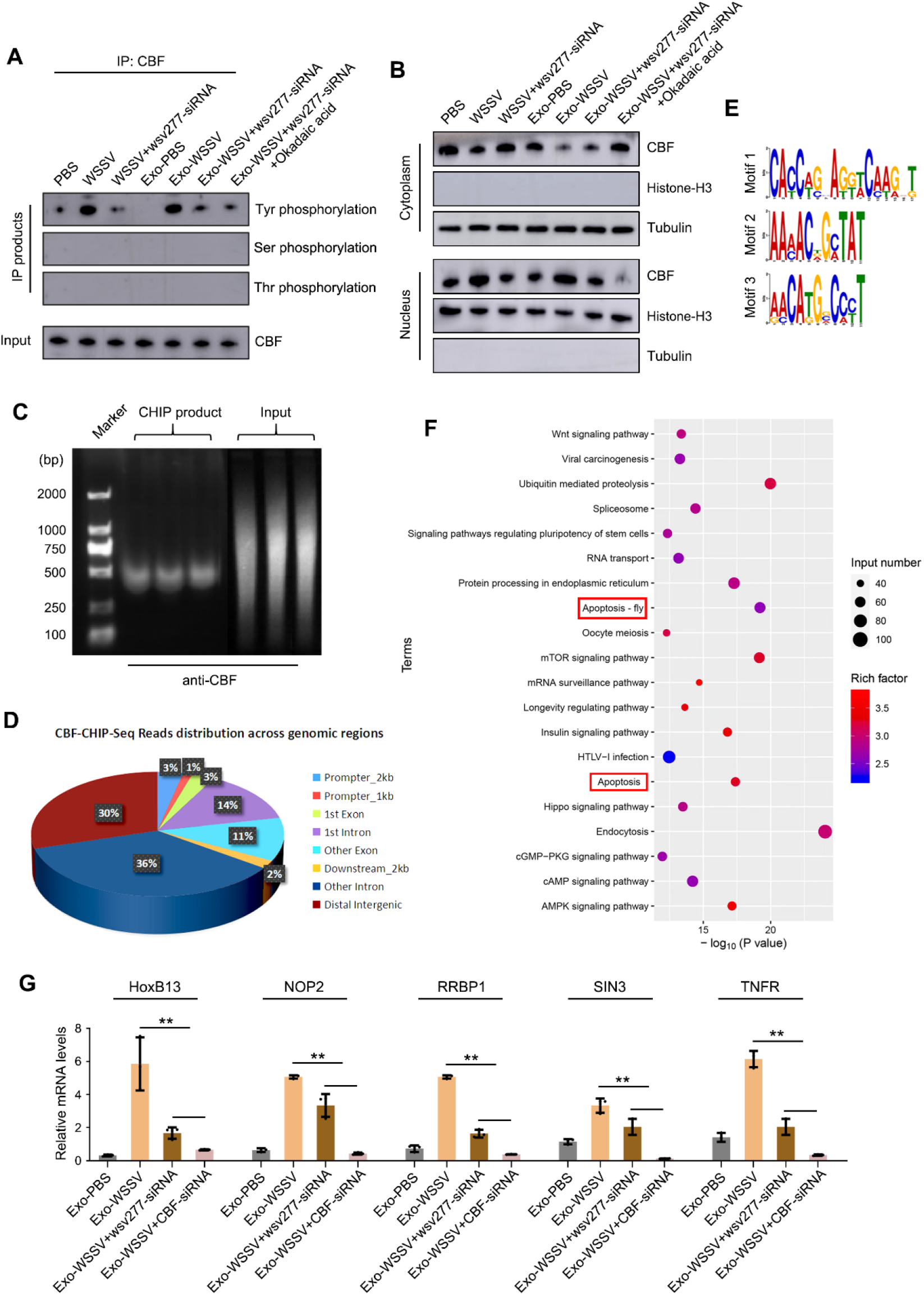
Mechanisms of apoptosis induced by CBF. **(A-B)** The influence of wsv277-packaged by exosomes on CBF phosphorylation and cellular distribution in hemocytes. Mud crabs were treated with WSSV, exosomes and wsv277-siRNA, followed by immunoprecipitated using CBF-antibody, then the IP products were subjected to phosphorylation detection at serine, threonine and tyrosine sites **(A)**; the cellular distribution of CBF was analyzed by detecting the protein levels of CBF in nucleus and cytoplasm, histone-H3 and tubulin were used for internal reference for nucleus and cytoplasm, respectively **(B)**. **(C)** Sample preparation for ChIP-seq with CBF -antibody. The product size was in the range of 200-500 bp. **(D)** CBF-ChIP-seq reads distribution across genomic regions. **(E)** The motifs of promoters bound to the CBF protein. **(F)** KEGG annotation of ChIP-seq data of CBF. The apoptosis-related pathways in the KEGG bubble chart were marked by red boxes. **(G)** The effects of wsv277 packaged by exosomes and CBF on the transcription of apoptosis-relevant genes. Five apoptosis-relevant genes (Including HoxB13, NOP2, RRBP1, SIN3 and TNFR) were selected from the data of ChIP-seq, β-actin was used as internal reference. The results are based on three parallel experiments and showed as mean values ± s.d. (*, *P*<0.05, **, *P*<0.01).

Next, we analyzed the downstream target genes of transcription factor CBF by ChIP-seq using anti-CBF IgG (Fig. 6C). The sequencing data of ChIP products showed that the reads distribution across genomic regions of CBF possess certain specificity (Fig. 6D), and the promoter sequences bound with CBF protein could be classified into 3 motifs (Fig. 6E). Then, KEGG analysis conducted on these genes revealed that many of CBF-regulated genes were involved in apoptosis pathway (Fig. 6F). To confirm these findings, five of CBF-regulated genes that have been reported to mediate apoptosis process were selected for further validation. The results showed that Exo-WSSV could promote the transcription of these apoptotic genes, and the positive effects were significantly inhibited when wsv277 or CBF was silenced (Fig. 6G). Taken together, the above data suggested that exosomes could mediate phosphorylation and nuclear translocation of CBF by wsv277, and then promote the transcription of apoptotic genes.

### wsv-277-EF-1α complex accelerate the synthesis of apoptotic proteins

To clarify the involvement of EF-1α in exosome-mediated apoptosis process, whether wsv277 could phosphorylate EF-1α needs further investigations. Thus, we analyzed the phosphorylation levels of EF-1α in mud crabs treated with exosomes and wsv277-siRNA, the results showed that wsv277 packaged by exosomes did not phosphorylate EF-1α (Fig. 7A). Meanwhile, the results of both western blot analysis and immune-fluorescence detection revealed that the cellular localization of EF-1α did not change after exosome treatment (Fig. 7B and S5A). Since EF-1α has been reported to be involved in protein synthesis during translation process^28^, therefore, we detected the levels of new-synthesized protein in the hemocytes by Click assay, the results showed that EF-1α was required for protein synthesis in mud crab (Fig. 7C, 7D and S5B). Moreover, through co-injection of exosomes and wsv277-siRNA, we found that Exo-WSSV was capable to accelerate protein synthesis by wsv277 (Fig. 7E, 7F and S5C).

**Fig 7.**
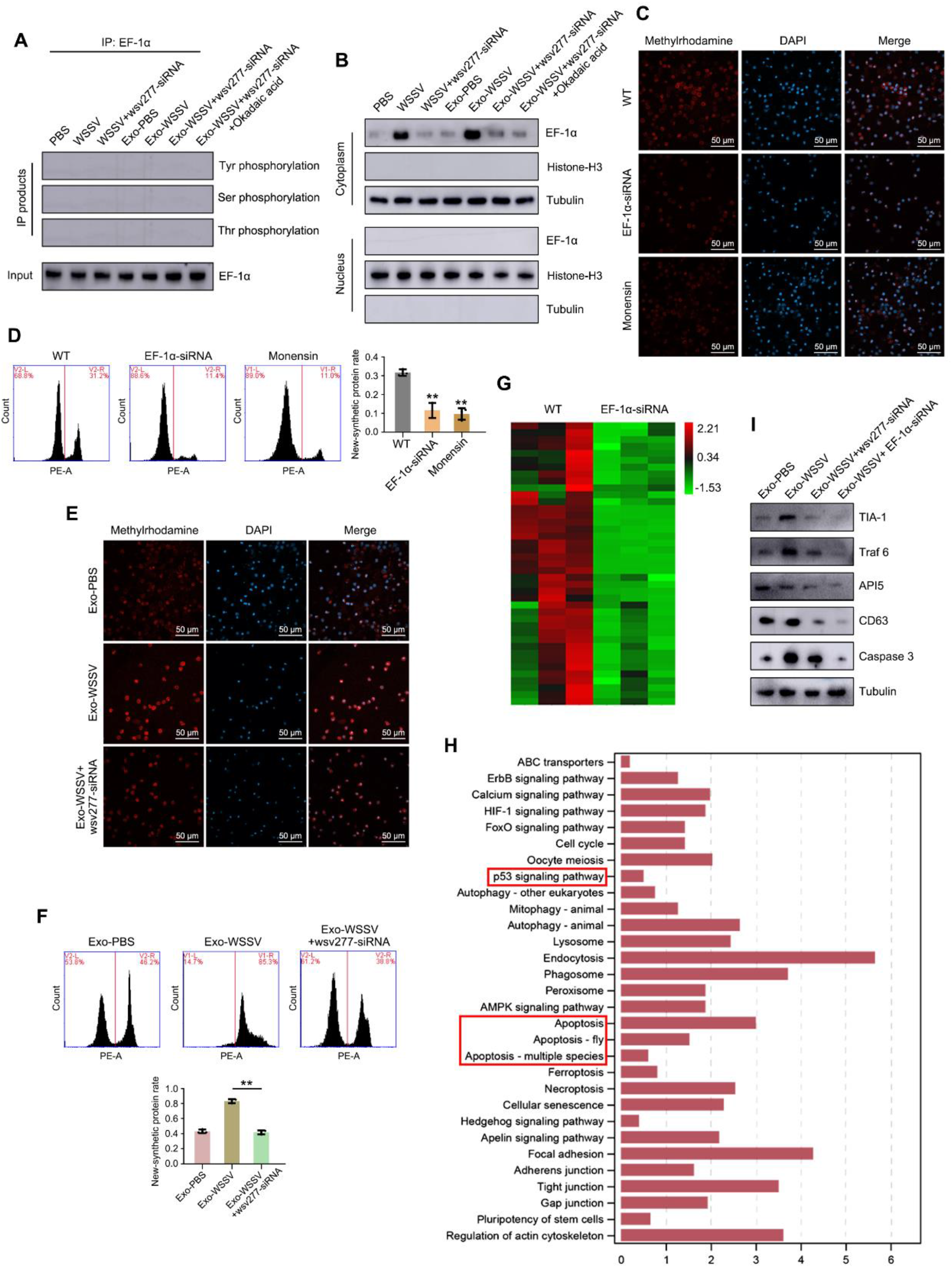
Mechanism of apoptosis induced by EF-1α. **(A-B)** The influence of wsv277-packaged by exosome on EF-1α phosphorylation and cellular distribution in hemocytes. Mud crabs were treated with WSSV, exosomes and wsv277-siRNA, followed by immunoprecipitated using EF-1α-antibody, then the IP products were subjected to phosphorylation detection at serine, threonine and tyrosine sites **(A)**; the cellular distribution of EF-1α was analyzed by detecting the protein levels of EF-1α in nucleus and cytoplasm, histone-H3 and tubulin were used for internal reference for nucleus and cytoplasm, respectively **(B)**. **(C-D)** The effects of EF-1α on protein synthesis in mud crab. Mud crabs were injected with EF-1α-siRNA for 48 h, then the levels of new-synthesized protein were detected by confocal microscopy (**C**) and flow cytometry **(D)** through performing Click assay, protein transport inhibitor Monensin was used as negative control. Scale bar, 50 μm. **(E-F)** The influence of wsv277 on exosome-mediated protein synthesis in mud crab. Mud crab were co-injected with exosomes and wsv277-siRNA, followed by new-synthesized protein detection through confocal microscopy (**E**) and flow cytometry **(F)**. **(G)** Proteomic analysis in hemocytes after EF-1α silencing. **(H)** KEGG annotation of proteome-seq data. The apoptosis-related pathways in the KEGG histogram were marked by red boxes. **(I)** The effects of wsv277 packaged by exosomes and EF-1α on protein synthesis of apoptosis-relevant genes. Five apoptosis-relevant genes (Including TIA-1, Traf6, API5, CD63 and Caspase 3) were selected from the data of proteome-seq, tubulin was used as internal reference. In all panels, data represent mean ± s.d. of triplicate assays (**, *P*<0.01).

The next issue is to identify the downstream proteins synthesized by EF-1α, thus, proteomic analysis was performed in the hemocytes after EF-1α knockdown. The most significantly downregulated proteins were subjected to KEGG analysis, and the results revealed that many of EF-1α-synthesized proteins were relevant to apoptosis pathway (Fig. 7G and 7H). To confirm the findings, five EF-1α-synthesized proteins that have been reported to mediate apoptosis process were selected for further validation, the results showed that Exo-WSSV could accelerate the synthesis of apoptotic proteins, and the acceleration effects was reduced after wsv277 or EF-1α silencing (Fig. 7I). Taken together, the above data suggested that wsv277 packaged by exosomes could cooperated with EF-1α, so as to accelerate the synthesis of apoptotic proteins.

In summary, the findings in this study indicated that during WSSV infection, viral mRNA wsv277 was specific encapsulated in exosomes secreted by host hemocytes, and then transmitted to the surrounding exosome-recipient cells, followed by translation into viral protein with phosphokinase activity, so as to promote the transcription and translation of apoptotic genes via interacting with CBF and EF-1α, respectively, which eventually suppressed the spread of virus infection by inducing apoptosis of the surrounding cells (Fig. 8A).

**Fig 8.**
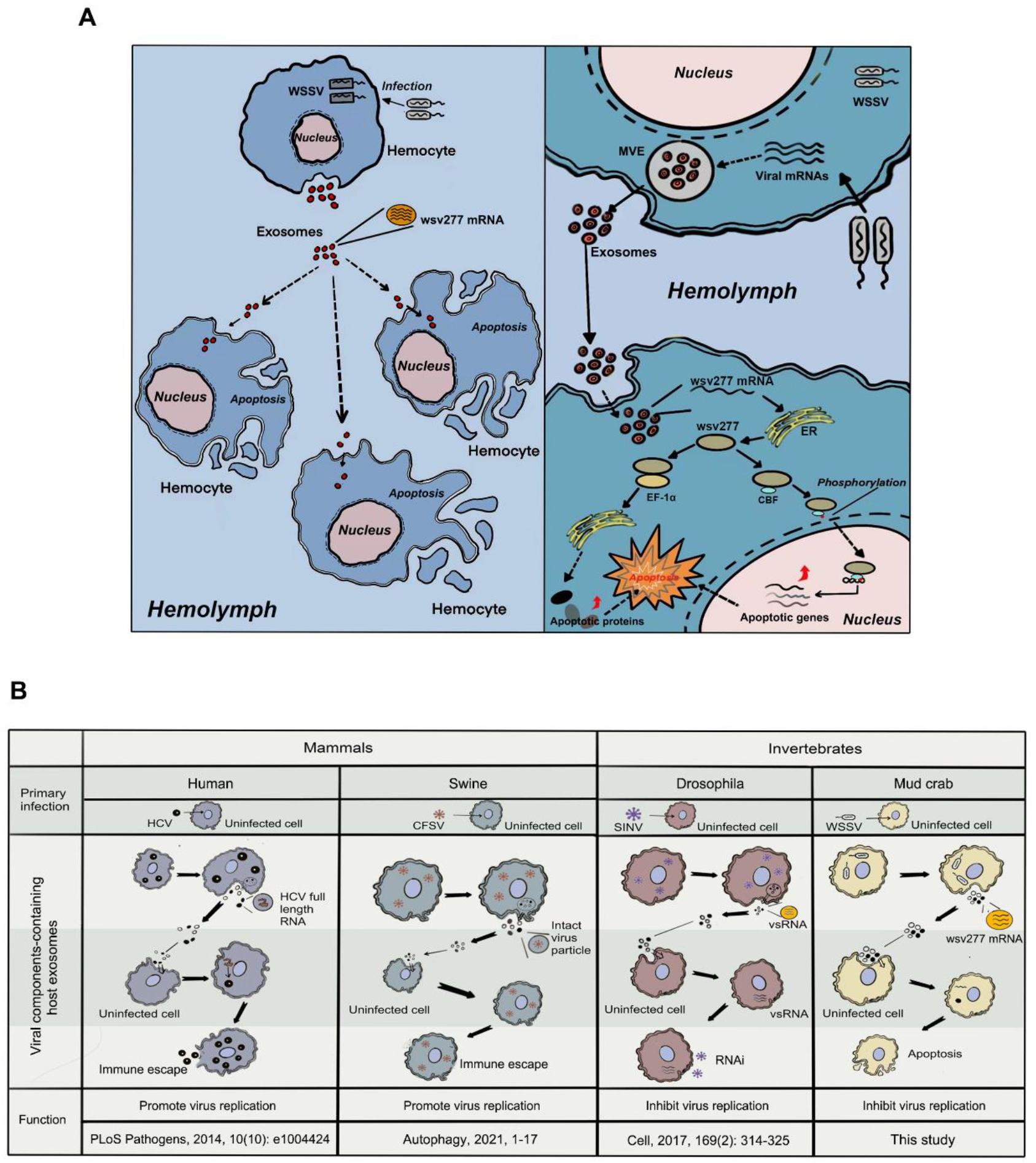
The proposed schematic diagram for exosome-mediated viral nucleic acid presentation and its role in innate immunity. **(A)** wsv277 packaged by exosomes contributes to anti-viral immune response in mud crab. During WSSV infection, viral mRNA wsv277 was encapsulated in exosomes secreted by host hemocytes, and then transmitted to the surrounding exosome-recipient cells, followed by translation into viral protein with phosphokinase activity, so as to promote the transcription and translation of apoptotic genes via interacting with CBF and EF-1α, respectively, which eventually suppressed the spread of virus infection by inducing apoptosis of the surrounding cells. **(B)** The role of exosome-mediated viral nucleic acid presentation in the innate immunity of vertebrates and invertebrates. In vertebrates, researches on human and swine have shown that virus could exploit exosomes to transmit viral particles or full-length genomic RNA and help to escape the host’s immune surveillance. While in invertebrates, results including this study indicate that host cells could drive exosomes to present viral substances, and further activate the immune state of surrounding exosome recipient cells to resist virus invasion.

## Discussion

As a novel approach of genetic exchange between cells, exosomes share similar biological sources with viruses, especially enveloped viruses^29,30^. Based on this, researchers have put forward the “Trojan exosome hypothesis” during virus infection^31^, the viral components, even infectious virus particles can be encapsulated and transmitted in exosomes to mediate non-receptor-dependent infection^32^. Recently, the involvement of exosomes during viral pathogenesis has been widely investigated^33^. For example, exosomes secreted from HIV-1 (Human immunodeficiency virus type-1)-infected macrophages contain HIV-1 virus particles and contribute to virus transfer to the uninfected cells^18^. Besides, during HCV (Hepatitis C virus) and EV71 (Enterovirus 71) infections, the full-length replication-competent viral RNA can be integrated into exosomes to mediate receptor-independent transmission of HCV to other susceptible cells^18,34^. Moreover, it was found that exosomes secreted by HSV (Herpes simplex virus)-infected cells could transport viral mRNAs microRNAs, so as to increase the susceptibility of un-infected cells to HSV^35^. At present, almost all studies relevant to viral components packaged by exosomes are reported in higher animals, and the virus type is RNA virus. Our results showed that viral mRNA wsv277-containing exosomes participated in the anti-viral immune response of mud crab, for the first time, we revealed that the components of DNA virus could be transmitted through the exosomes of lower invertebrates, which strongly demonstrated the diversity of exosome-mediated viral immunity and its universality in animals.

Apoptosis is one type of crucial cellular immune response that enable the hosts to protect themselves against pathogenic microbes^36^, the correlation between apoptosis and virus infection and its regulatory mechanism have always been the hot topics in the field of innate immunity^37^ Generally, as a multi-factor process, apoptosis could be activated during virus infection to eliminate invading virus particles, and serves as an effective defense tool to maintain homeostasis of the host^38^. Another situation is that the virus-encoded genes have evolved the ability to inhibit or delay the apoptosis of infected cells to ensure virus life cycle, and it is a survival mechanism by which the virus escapes the host’s innate immune defense^39,40^. Interestingly, some researchers found that the hemocytes of shrimp infected by WSSV particles did not undergone apoptosis, but the uninfected surrounding hemocytes showed severe apoptosis^41,42^ So far, there is no reasonable explanation for this finding. To explain the phenomenon, it is essential to clarify how the virus-infected host cells transmit apoptotic signals to the surrounding cells. Our study showed that exosomes secreted from WSSV-infected mud crab hemocytes contain viral nucleic acid wsv277 and could be internalized into exosome-recipient cells, and further activated the transcription and translation of apoptotic genes by interacting with CBF and EF-1α, so as to prevent the spread of virus infection through inducing apoptosis of the surrounding cells. Therefore, for the first time, we elucidated the mechanism of apoptotic signal transduction between cells from the perspective of exosome, and revealed a novel strategy for the host to cope with viral infection.

The innate immune system of animals is evolved during the long-term natural selection, and its function and physiological significance have been widely recognized^43^. With the continuous evolution of the vertebrates, lymphoid cells (B cells and T cells) and specialized lymphoid organs appeared, and finally developed into acquired immune system mainly based on specific humoral immunity and cellular immunity^44^. It has been found that the CD4^+^ T cell-dependent anti-virus antibody production was crucial for clearing viral particles following acute vaccinia virus infection^45^. Besides, the early induction of functional SARS-CoV-2 specific T cells were tightly relevant to rapid viral clearance and mild symptoms in COVID-19 patients^46^. These studies indicated that apart from innate immunity, acquired immunity also serves as essential strategy for vertebrate to defense against viral infection, and has the characteristics of specificity and memorability^47^. In this case, whether the role of innate immune system is weakened or even degraded in vertebrates has been widely concerned by immunologists^48^. In our opinion, this problem could be clarified to some extent from the perspective of exosomes during host-virus interactions. In vertebrates, all relevant researches have shown that virus could exploit exosomes to transmit viral particles or full-length genomic RNA and help to escape the host’s immune surveillance^30,34,49^ While in invertebrates, results including this study indicate that host cells could drive exosomes to present viral substances, and further activate the immune state of surrounding exosome recipient cells to resist virus invasion^25^ (Fig. 8B). Therefore, since exosome is an important innate immune regulation approach, we hypothesize that the innate immune system of vertebrates is weakened or degenerated, allowing exosomes to be employed by virus as the vehicle for immune escape.

## Materials and Methods

### Ethics statement

The mud crabs used were taken from Niutianyang aquaculture farm (Shantou, China). No specific permits were required since mud crab was not belonged to endangered or protected species. The animals were processed according to “the Regulations for the Administration of Affairs Concerning Experimental Animals” established by the Guangdong Provincial Department of Science and Technology on the Use and Care of Animals.

### WSSV challenge and detection

Healthy mud crabs, approximately 50 g each, were domesticated in a thermostatic culture tank at 25°C at 10‰ salt seawater for 72 hours before WSSV challenge. PCR analysis was performed via WSSV-specific primer (5’-TATTGTCTCTCCTGAC-GTAC-3’ and 5’-CACATTCTTCACGAGTCTAC-3’) to ensure the mud crabs used were WSSV-free before treatments. Then, 200 μL of WSSV suspension (1×10^6^ particles/mL) was injected into each crab, hemolymph and tissues of 9 randomly chosen crabs per group was collected for later use at different time post-injection. After that, WSSV copies was detected by qPCR analysis using Premix Ex Taq (Probe qPCR) (Takara, Japan) with WSSV primers (5’-TTGGTTTCATGCCCGAGATT-3’, 5’-CCTTGGTCAGCCCCTTGA-3’) and a TaqMan probe (5’-FAM-TGCTGCCGTCTCC AA-TAMRA-3’).

### Isolation and identification of exosomes

In order to separate exosomes from the hemolymph of mud crabs, hemolymph was resuspended and centrifuged at 8,000 ×*g*, 4°C for 30 min, followed by centrifugation of the supernatant at 20,000 ×*g*, 4°C for 1 h, finally the above supernatant was centrifuged at 130,000 ×*g*, 4°C for 2 h. After that, the collected sediment was suspended with 2 mL of 0.95 M sucrose solution, and then 2 mL of 2.0 M sucrose solution, 2 mL of 1.3 M sucrose solution and 2 mL of 0.95 M sucrose solution were sequentially added before being centrifuged at 4 °C, 200,000 ×*g* for 16 h. The solution between 1.3 M and 0.95 M sucrose was filtered with 0.22 μm filters and dialyzed with an 8-15 kDa dialysis bag for 3 h. The obtained exosomes were observed under Philips CM120 BioTwin transmission electron microscope (FEI Company, USA) and quantified with Nano-Sight NS300 system (Malvern Instruments, UK).

### RNA interference assay

Based on the sequence of wsv001, wsv091, wsv271, wsv277, wsv447, CBF and EF-1α (GenBank accession numbers are AY048547.1, JX515788.1, MH663976.1, MN840357.1, KT995472.1, ON464159 and ON464160), the siRNAs were designed as follows: wsv001-siRNA (5’-CAGGGCCGTGTATGAAGCGTCAATT-3’), wsv094 - siRNA (5’-CCAGTTCCAGTTATCAACATCAAAT-3’), wsv271-siRNA (5’-GCTCA ACATTTAGTGGACATGACAA-3’), wsv277-siRNA (5’-TCGAAAGGACCGGAGAGCCTCTTAA-3’), wsv477-siRNA (5’-CCAGTTCCAGTTATCAACATCAAAT-3’), CBF-siRNA (5’-GGGTCTCCCTGTATGTGATAGCAGT-3’) and EF-1α-siRNA (5’-CAGGGCCGTGTATGAAGCGTCAATT-3’). *In vitro* Transcription T7 Kit (Takara, China) was used to synthesize specific siRNAs targeting the above genes according to the user’s instructions. Then, 1 μg of siRNAs were injected into per gram of crab. At different time post injection, 3 mud crabs for each group were randomly selected and stored for later use.

### Quantification of mRNA with qPCR

Total RNA of hemocytes was extracted by RNAiso Plus (Takara, Japan), followed by first-strand cDNA synthesis through PrimeScript RT Reagent Kit (Takara, Japan). Primers used were listed as follows: wsv001-F (5’-ATCGGGTGTTTTTGTG G-3’), wsv001-R (5’-GGTGTTTTTGTGGGCTA-3’), wsv091-F (5’-GATGTTCGCC CTTATGA-3’), wsv091-R (5’-TTGAGCGGAAGTTGATG-3’), wsv271-F (5’-TTTT GTTGGCAAGATTG-3’), wsv271-R (5’-CGATAGATATAGCCGTTTT-3’), wsv277 - F (5’-TTTGGCTCAGGAGAGAG-3’), wsv277-R (5’-TTTTGTTGGCAAGATTG-3’), wsv447-F (5’-TACCCAGGATGCGTATG-3’), wsv447-R (5’-ACCCCGTTCAA CAGAAT-3’), CBF-F (5’-GCTGGACTGTGTTCCTAAACC-3’), CBF-R (5’-GCTG GACTGTGTTCCTAAACC-3’), EF-1α-F (5’-GCTGGACTGTGTTCCTAAACC-3’) and EF-1α-R (5’-CAGCGAACTTGCAGGCGATATG-3’), while primers β-actin (F, 5’-GCGGCAGTGGTCATCTCCT-3’ and R, 5’-GCCCTTCCTCACGCTATCCT-3’) was used the internal control. Relative fold change of mRNA levels was determined using the 2^-ΔΔCt^ algorithm.

### Detection of apoptotic activity

The apoptosis rate of crab hemocytes was evaluated through detecting Annexin V, mitochondrial membrane potential and Caspase-3 activity. Annexin V was assessed by Annexin V FITC apoptosis assay kit (BD Pharmingen, USA) and measured by Accuri C6 Plus flow cytometry (BD Biosciences, USA). Mitochondrial membrane potential was analyzed by mitochondrial membrane potential test kit JC-1 (Beyotime, China) and observed under a confocal microscopy (ZEISS, Germany), Caspase-3 activity was measured by Caspase-3 activity detection kit (Beyotime, China) and tested by a Flex Station II microplate reader (Molecular Devices, USA).

### Protein structural analysis and kinase activity detection

The secondary structure of wsv277 protein was predicted by (http://smart.embl-heidelberg.de). The tertiary structure of was predicted by swiss-Model. wsv277 kinase site was analyzed by (https://www.phosphosite.org). Kinase-Lumi Enhanced Chemiluminescence Kinase Activity Detection Kit (Beyotime, China) was used to measure the kinase activity of wsv277 protein.

### Cell culture, transfection, and coIP assays

The *Drosophila* Schneider 2 (S2) cells were cultured in Schneider’s *Drosophila* Medium (with 10% (w/v) FBS) (Invitrogen, USA) at 28 °C. The constructed plasmids including pAc5.1A/V5-His-wsv277, pAc5.1A/V5-His-HA-CBF, pAc5.1A/V5-His-HA-EF-1α, pAc5.1A/V5-His-CBF and pAc5.1A/V5-His-EF-1α were transfected into S2 cells using Cellfectin II Reagent (Invitrogen, USA), respectively, pAc5.1A/V5-His and pAc5.1A/V5-His-HA were used as negative control. 48 h after co-transfection, western blot was conducted to analysis the interaction between wsv271, CBF and EF-1α.

### Fluorescence *in situ* hybridization

Crab hemocytes were inoculated on confocal dishes treated with 10% polyresistant acid and fixed with 4% paraformaldehyde for 15 min at room temperature, and then dehydrated with 70% ethanol and incubated at 4 °C overnight, followed by incubation with hybrid liquid [1× SSC (15 mM sodium citrate, 150 mM sodium chloride, pH 7.5) and 10% (w/v) dextran sulfate] for 0.5 h, subsequently, wsv277 fluorescent probe (5’-FAM-TTGGATGAAGAACTAGTTTCTT-3’) was added to the hybrid solution with final concentration of 500 mM. After washed with PBS for 3 times and stained with DAPI (4’,6-diamidino-2-phenylindole) (50 ng/mL) (Sigma, USA) for 5 min, the slips were observed using a CarlZeiss LSM710 system (Carl Zeiss, Germany).

### Chromatin Immunoprecipitation assay

The ChIP Assay Kit (Beyotime, China) was used in this experiment. Firstly, crab hemocytes were ultrasonically broken and cross-linked by formaldehyde, then, the genomic DNA was sheared into 200-1000 bp through 270 kW ultrasound. Then, the samples were mixed with Dilution Buffer containing 1mM of PMSF, followed by incubated with Protein A+G (Agarose/Salmon Sperm DNA) for 30 min at 4 °C. After that, CBF antibody was added and mixed overnight at 4 °C, finally, the mixtures were washed with elution buffer for twice, and the flow through fluids were collected and used for agarose gel electrophoresis. The samples were sequenced and analyzed by Wuhan Kang Testing Company (China), and the raw data have been uploaded to NCBI public database (Accession number: PRJNA838265).

### New-synthetic protein detection

Click-IT AHA (L-azido-homoalanine) was used to determine the levels of new-synthetic proteins in crab hemocytes. Firstly, 25 mg/mL of Click-IT AHA was added into methionine-free cell culture medium to incubate with hemocytes for 1h, followed by infiltrated with PBS (0.25% Triton X-100) for 15 min. Subsequently, PBS containing 25 μg/mL of 5-carboxyltetramethylrhodamine-azide was added and incubated for 15 min at room temperature. After incubated with Hoechst or DAPI at room temperature for 15 min, the intensity of red fluorescence in hemocytes was measured by confocal microscopy (Carl Zeiss, Germany), microplate reader (Molecular Devices, USA) and flow cytometry (BD Biosciences, USA).

### Transcriptome and proteome sequencing

The transcriptome of exosomes in mud crab were sequenced by Beijing Biomarker Biotechnology Co., Ltd. (China), the processing groups include PBS group and WSSV group, and the original data have been uploaded to NCBI public database (Accession number: PRJNA837743). The transcriptome of hemocytes treated with Exosome-PBS, Exosome-WSSV and Exosomes-WSSV + wsv277-siRNA were sequenced by Kidio Biotechnology Co., Ltd. (Guangzhou, China), the raw data have been uploaded to NCBI public database (Accession number: PRJNA837866). The proteome of hemocytes were sequenced by Kidio Biotechnology Co., Ltd. (Guangzhou, China), and the processing groups include WT (Wild type) group and EF-1α-siRNA group, the raw data have been uploaded to iProX public database (Accession number: IPX0004462001).

### Statistical analysis

One-way ANOVA analysis was performed for all data through using Origin Pro 8.0 software, *P*<0.05 means the difference was statistically significant. All assays were biologically repeated for three times.

## Acknowledgments

This study was financially supported by National Natural Science Foundation of China (32173006, 42076125), 2020 Li Ka Shing Foundation Cross-Disciplinary Research Grant (2020LKSFG01E) and Key Special Project for Introduced Talents Team of Southern Marine Science and Engineering Guangdong Laboratory (Guangzhou) (GML2019ZD0606). The funders had no role in study design, data collection and analysis, decision to publish, or preparation of the manuscript.

## Author contributions

YG, HH, XSZ, WQW, MZ and TNT performed the experiments, YG and SKL designed the experiments and analysed the data, HYM and YLZ provided technical supports, YG and SKL wrote the manuscript. All authors read and approved the contents of the manuscript and its publication.

## SUPPLEMENTAL DATA

**Fig S1.**
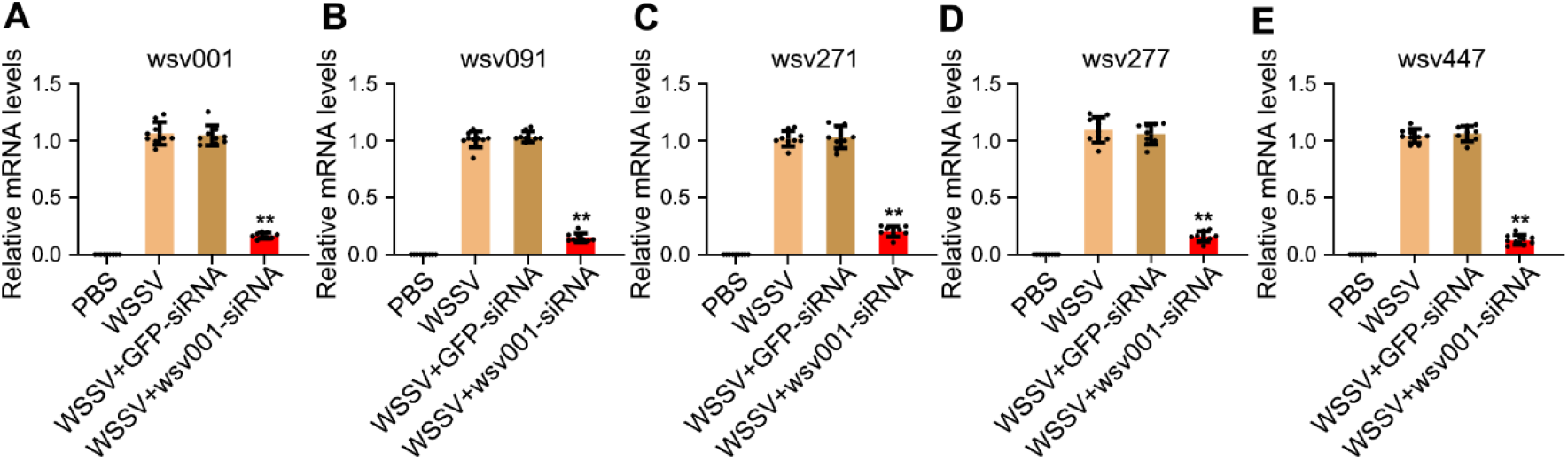
RNAi efficiency detection of five viral nucleic acids packaged by exosome. **(A-E)** Mud crabs were co-injected with siRNAs targeting wsv001, wsv091, wsv271, wsv277 and wsv447 for 24 h, then the mRNA expressions of these viral genes were detected by qPCR, β-actin was used as internal reference. Data were the mean ± s.d. of three replicated experiments (**, *P*<0.01).

**Fig S2.**
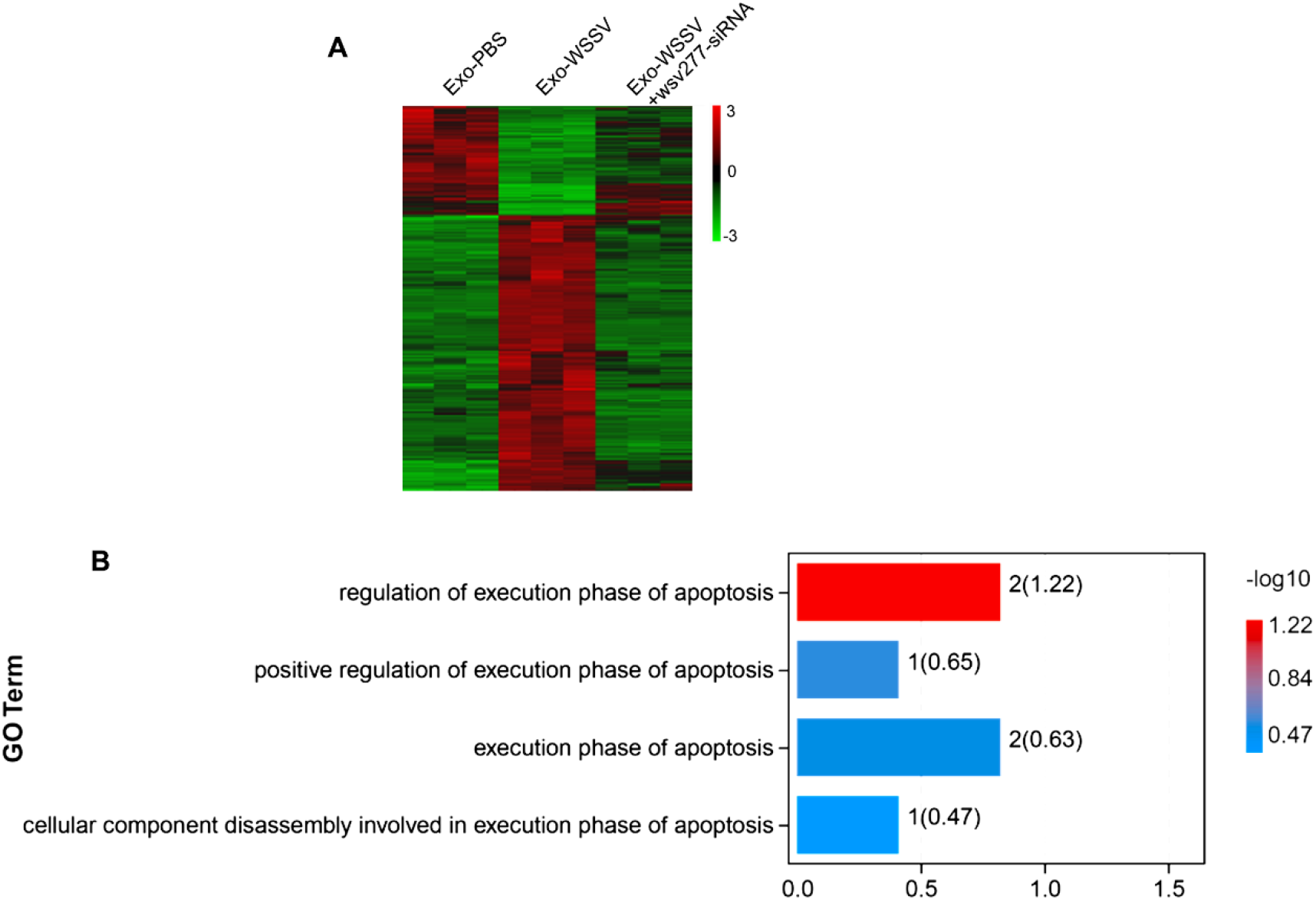
**(A)** Transcriptome analysis of mud crab challenged with exosomes and wsv277-siRNA, 428 differentially expressed genes that altered in Exo-WSSV group compared with Exo-PBS, and with no significant difference when wsv277 was silenced, were shown on the heat map. **(B)** Apoptotic functional genes screened by GO analysis were presented in histogram.

**Fig S3.**
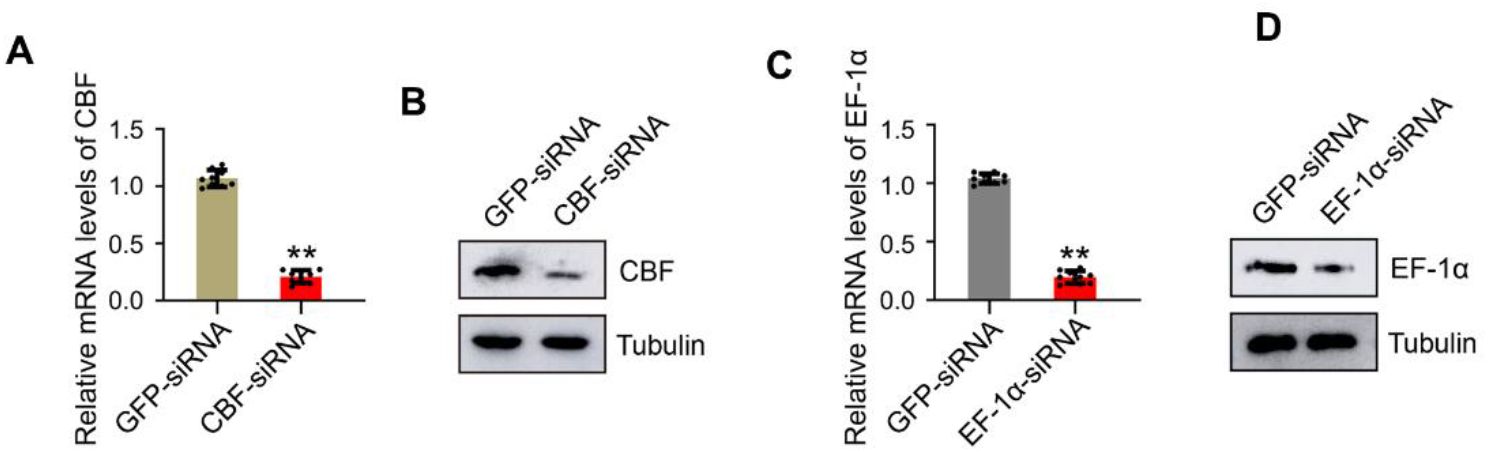
RNAi efficiency detection of CBF and EF-1α. **(A-D)** Mud crabs were injected with siRNAs targeting CBF or EF-1α for 24 h, then both the mRNA and protein levels of CBF **(A-B)** and EF-1α **(C-D)** were detected by Western blot and qPCR, respectively. Data are based on three parallel experiments and shown as mean values ± s.d. (*, *P* < 0.05, **, *P* < 0.01).

**Fig S4.**
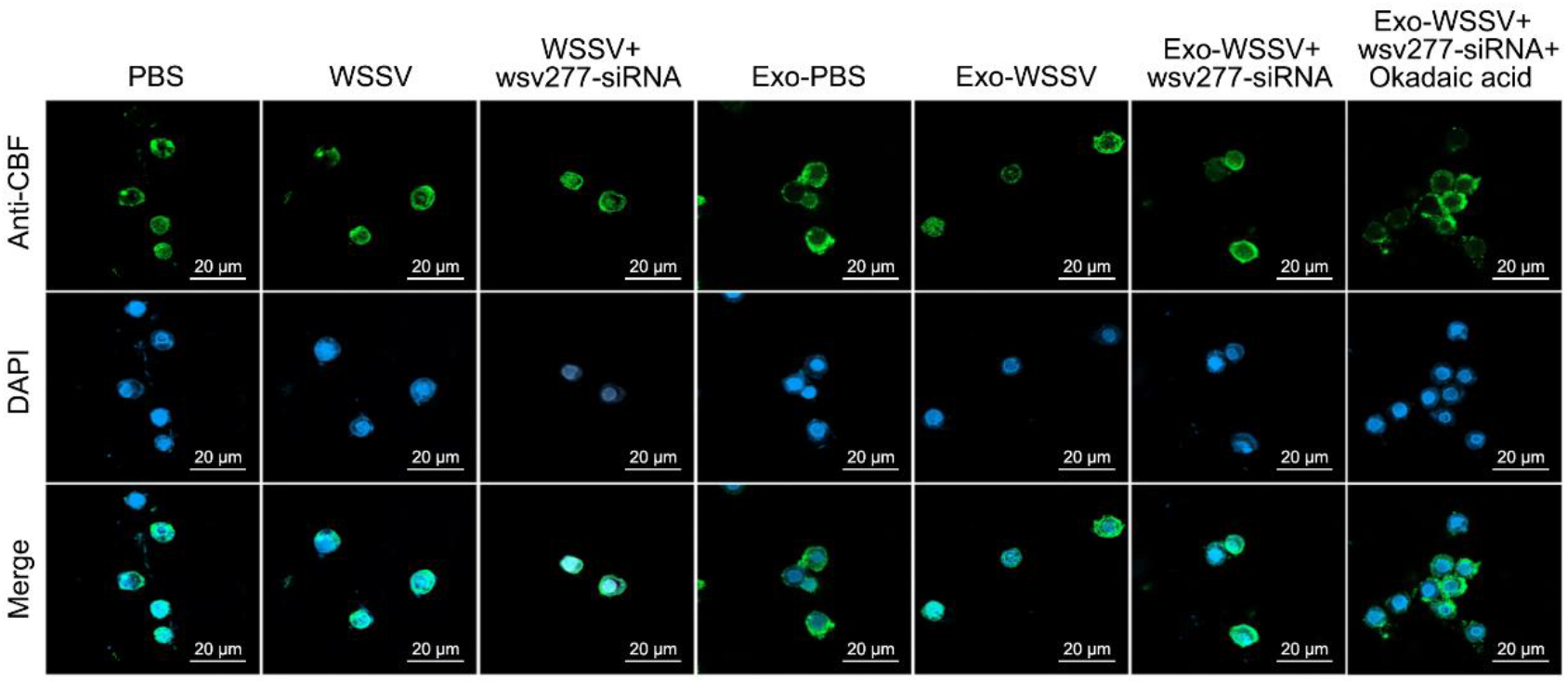
The cellular localization of CBF in the hemocytes of mud crab. Mud crabs were treated with WSSV, exosomes and wsv277-siRNA, then the localization of CBF was determined using immunofluorescence assay with CBF antibody, DAPI was used to stain nucleus. Scale bar, 20μm.

**Fig S5.**
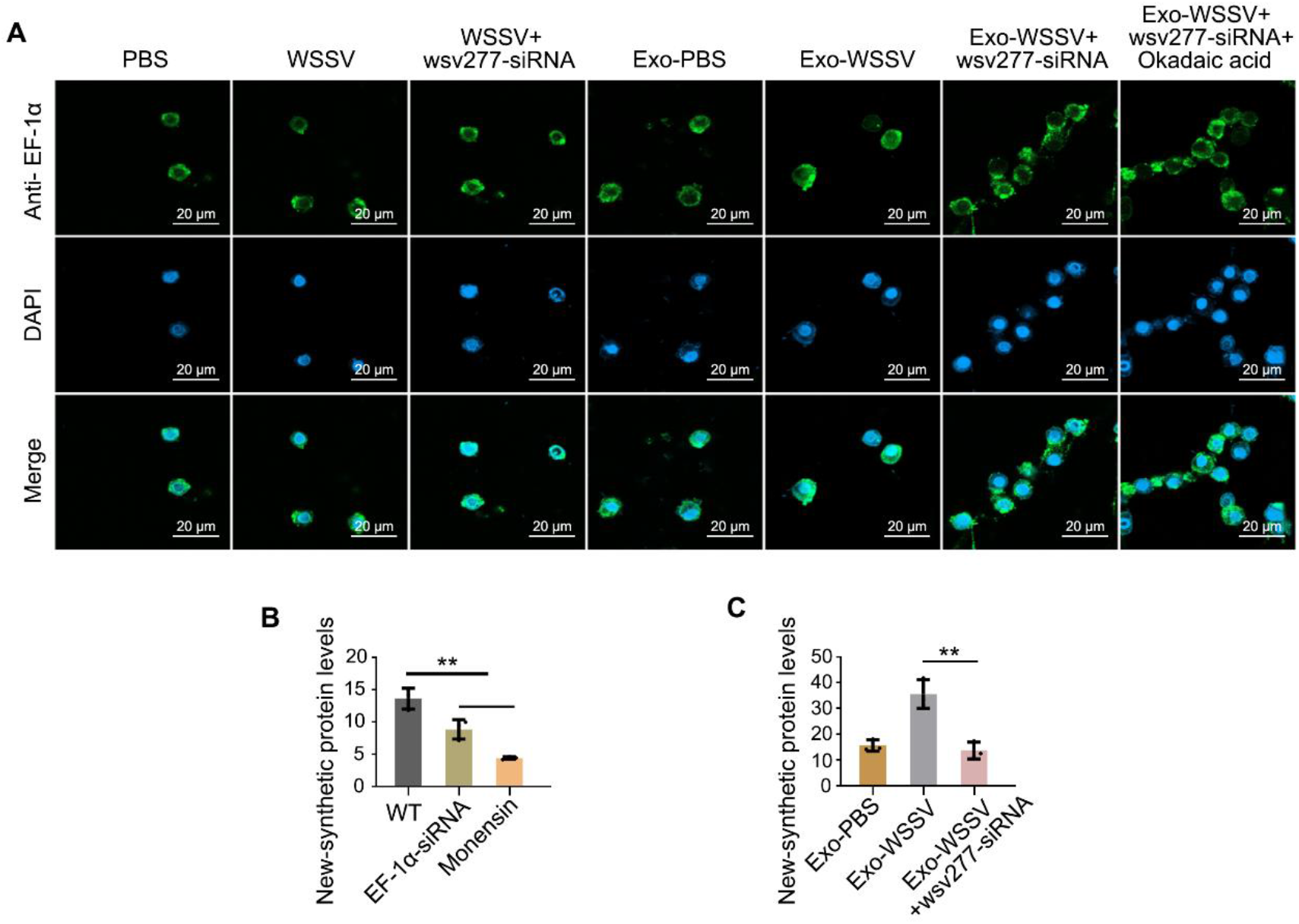
**(A)** The cellular localization of EF-1α in the hemocytes of mud crab. Mud crabs were treated with WSSV, exosomes and wsv277-siRNA, then the localization of EF-1α was determined using immunofluorescence assay with EF-1α antibody, DAPI was used to stain nucleus. Scale bar, 20μm. **(B)** The effects of EF-1α on protein synthesis. Mud crabs were injected with EF-1α-siRNA for 48 h, then the levels of new-synthesized protein were detected with a microplate reader. **(C)** The influence of wsv277 on exosome-mediated protein synthesis. After co-treated with exosomes and wsv277-siRNA, mud crabs were subjected to new-synthesized protein detection with a microplate reader. Data shown represent the mean ± s.d. for triplicate assays (**, *P* < 0.01).

